# The rise and fall of feminizing *Wolbachia* in a butterfly

**DOI:** 10.64898/2026.10.19.760017

**Authors:** Kei Ishida, Akira Matsubara, Daisuke Kageyama, Mai Miyata

## Abstract

In sexually reproducing animals, sex ratios generally evolve toward parity^1^ but can be distorted by selfish genetic elements, including sex chromosome-linked meiotic drivers and cytoplasmically inherited symbionts such as *Wolbachia*^2^. We previously documented the rapid spread of the feminizing *Wolbachia* strain *w*Fem in an island population of the butterfly *Eurema hecabe*^3^. Within just four years, the proportion of females increased from approximately 50% in 2015–2018 to 93.1% in 2022. Here, continued monitoring of the same population revealed an equally striking reversal: the sex ratio returned to near parity in 2024–2025, accompanied by a sharp decline in the frequency of *w*Fem. These observations suggest that host–symbiont conflict can drive major shifts in population sex ratio over remarkably short timescales.

## Main text

Throughout our 2015–2022 surveys, all examined *w*Fem-positive females (hereafter CF females) produced exclusively female offspring, whereas *w*Fem-negative females (C females) produced offspring with an approximately 1:1 sex ratio. If feminization imposes strong selection on the host, however, host genetic variants that counteract feminization may be favored and spread, as expected under an evolutionary arms race^4^. Rapid evolution of host suppression has previously been documented in *Hypolimnas* butterflies^5,6^ and *Mallada* lacewings^7^ in response to male-killing symbionts. Remarkably, in *E. hecabe*, we captured both the rapid rise of feminization and its subsequent decline within a single decade.

In 2023, the sex ratio of individuals collected from the focal population remained strongly female-biased, with females accounting for 82.3% of the individuals (Figure 1A). CF females accounted for 85.3% of the females collected that year, but in striking contrast to previous years, they were highly heterogeneous in terms of the offspring sex ratio. Of the 20 CF females, 10 produced exclusively female offspring, seven produced exclusively male offspring, and three produced both males and females (Figure 1B). In 2024, the proportion of CF females declined only slightly to 80.6%, whereas the sex ratio shifted sharply toward parity, with females accounting for 52.2% of the collected individuals (Figure 1A). CF females continued to show marked heterogeneity in offspring sex ratio (Figure 1B). By 2025, the change had progressed further: the proportion of CF females dramatically dropped to 28.3%, whereas females accounted for only 36.0% of the collected individuals (Figure 1A).

**Figure 1.**
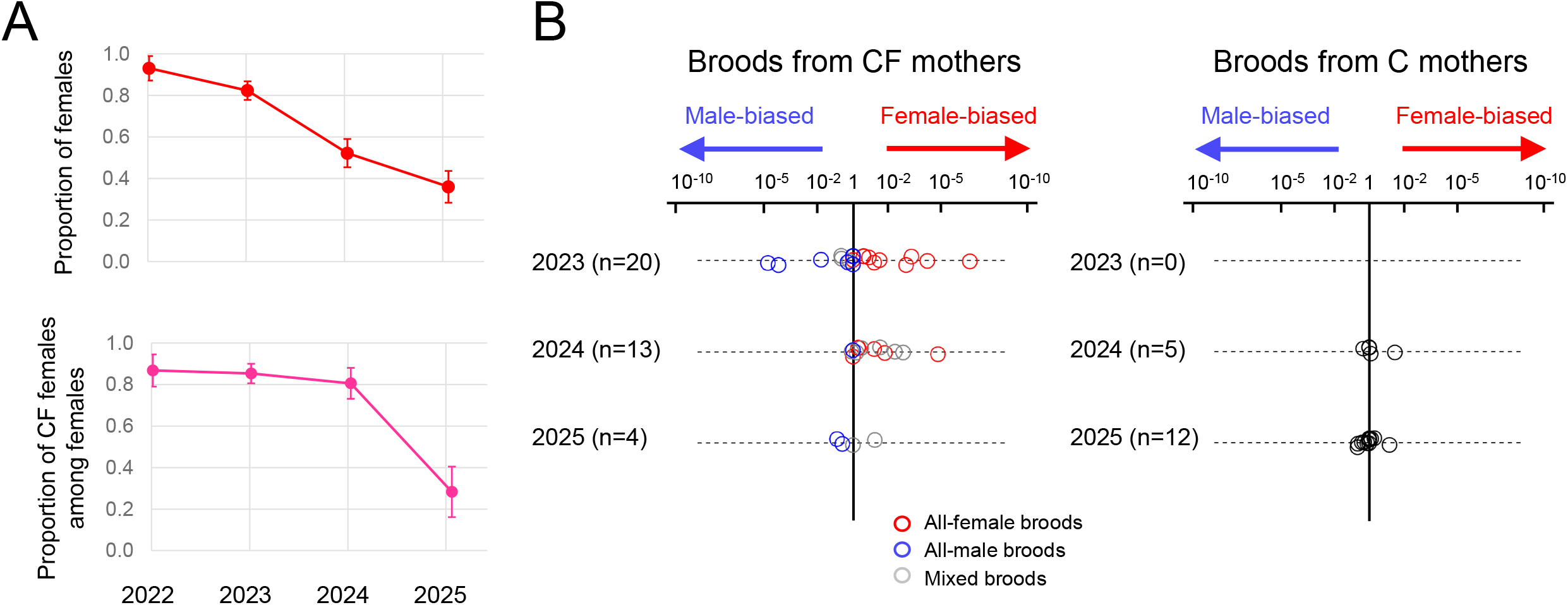
*Eurema hecabe* adults collected on Ishigaki Island and their offspring. (A) Proportions of females among the collected adults (top) and proportions of *w*Fem-positive (CF) females among the females (bottom), both with 95% confidence intervals. (B) Offspring sex ratios of 54 field-collected mothers. Each circle represents a single brood. Broods produced by CF mothers are shown on the left, and those produced by *w*Fem-negative (C) mothers on the right. P values on the x-axis were calculated using two-sided exact binomial tests against an expected 1:1 sex ratio. The direction of deviation is indicated by plotting male- and female-biased broods to the left and right of the central axis, respectively.

The mechanism underlying this rapid reversal remains unknown, but offspring infection patterns provide an important clue. All daughters examined from CF mothers (n = 31), whether derived from all-female or mixed broods, carried *w*Fem. In contrast, all sons examined (n = 15), whether derived from the all-male or mixed broods, lacked *w*Fem (Table S1). Thus, transmission of *w*Fem was associated with offspring sex: daughters inherited *w*Fem, whereas sons did not.

This pattern resembles that observed in the sister species *Eurema mandarina*, in which antibiotic treatment of CF females disrupted feminization^8^. In the present population, however, the *Wolbachia* density did not significantly differ among CF females producing broods with different sex ratios (Figure S1). One possibility is that host genetic variation affects *w*Fem specifically in the germline or during transmission to the offspring, rather than simply reducing its overall abundance in the mother. This effect could generate all-female, mixed, and all-male broods if the efficiency of suppression varies among females. More generally, the tight association between loss of *w*Fem and male development raises the possibility that the suppression of feminization acts, directly or indirectly, by destabilizing maternal transmission of *w*Fem. However, the detection of a very small number of *w*Fem-positive males among the collected males (7 of 245 males) indicates that complete loss of *w*Fem is not always required for male development. These rare males may have inherited *w*Fem at a density sufficient for transmission but below that required to induce feminization, suggesting that the threshold for feminization is higher than that for maternal transmission (Supplemental discussion).

Taken together, these consecutive observations captured a remarkable bidirectional shift in the sex ratio of *E. hecabe*: from parity to an extreme female bias and then back toward parity, all within a decade spanning 2015 to 2025. Over the same period, the frequency of *w*Fem increased from undetectable levels to 80.6% and subsequently declined sharply to 28.3%. Notably, *w*Fem had previously been detected at a frequency of 8% in this population in 2008^9^, raising the possibility that the recent rise and fall reflects recurrent dynamics rather than a singular evolutionary event in this population.

We hypothesize that the extreme scarcity of males during the period of strong female bias imposed intense selection favoring host genotypes that restore male production. If such suppression simultaneously compromises transmission of *w*Fem, a selection for the rare sex could also drive a subsequent decline in the feminizing symbiont itself. This differs from several male-killing systems, in which suppression of male killing restores male survival without necessarily impairing maternal transmission of the symbiont. Consequently, suppressors of male killing can spread while the symbiont remains common in the population^10^.

The exceptionally rapid turnover observed in *E. hecabe* may therefore reflect a distinctive feature of feminization systems: the restoration of male development appears to be coupled to failure of transmission of the feminizing symbiont. Such coupling could allow host suppression not only to restore the population sex ratio but also to undermine transmission of the selfish element that created the sex-ratio distortion. More generally, host–distorter arms races may proceed particularly rapidly when suppression directly reduces transmission of the distorter itself.

## Acknowledgements

We thank Masashi Nomura, Yuhei Hashimoto, and Rinka Kimoto for collecting the butterflies and Waka Minami for assistance with insect rearing. This work was supported by a JSPS KAKENHI grant (25K00108) and Sumitomo Foundation for basic Sciences to M.M. and a JSPS KAKENHI grant (23K26922 and 24H02293) to D.K.

## Author contributions

K.I., D.K., and M.M. designed the study. K.I., A.M., and M.M. conducted field sampling, rearing experiments, and molecular experiments. K.I. and M.M. analyzed the data. K.I., D.K., and M.M. wrote the manuscript. All authors read and approved the final manuscript.

## Declaration of generative AI and AI-assisted technologies in the manuscript preparation process

During the preparation of this work, the authors used ChatGPT to assist with grammar and style. The authors reviewed all AI-assisted edits and take full responsibility for the final content.

## Supplemental information

Table S1

Figure S1

Supplemental experimental procedures

Supplemental discussion

## REFERENCES

1 Fisher, R. (1930). The Genetic Theory of Natural Selection. (Clarendon Press).

2 Beukeboom, L. W., and Perrin, N. (2014). The Evolution of Sex Determination. (Oxford University Press). 10.1093/acprof:oso/9780199657148.001.0001.

3 Miyata, M., Nomura, M., and Kageyama, D. (2024). Rapid spread of a vertically transmitted symbiont induces drastic shifts in butterfly sex ratio. Curr. Biol. 34, R490–R492. 10.1016/j.cub.2024.04.027.

4 Hamilton, W. D. (1967). Extraordinary sex ratios. A sex-ratio theory for sex linkage and inbreeding has new implications in cytogenetics and entomology. Science 156, 477–488. 10.1126/science.156.3774.477.

5 Charlat, S., Hornett, E. A., Fullard, J. H., Davies, N., Roderick, G. K., Wedell, N., and Hurst, G. D. D. (2007). Extraordinary flux in sex ratio. Science 317, 214–214. 10.1126/science.1143369.

6 Hornett, E. A., Charlat, S., Duplouy, A. M. R., Davies, N., Roderick, G. K., Wedell, N., and Hurst, G. D. D. (2006). Evolution of male-killer suppression in a natural population. PLOS Biol. 4, e283. 10.1371/journal.pbio.0040283.

7 Hayashi, M., Nomura, M., and Kageyama, D. (2018). Rapid comeback of males: evolution of male-killer suppression in a green lacewing population. Proc. Biol. Sci. 285, 20180369. 10.1098/rspb.2018.0369.

8 Kageyama, D., Ohno, M., Sasaki, T., Yoshido, A., Konagaya, T., Jouraku, A., Kuwazaki, S., Kanamori, H., Katayose, Y., Narita, S., et al. (2017). Feminizing Wolbachia endosymbiont disrupts maternal sex chromosome inheritance in a butterfly species, Evol. Lett. 1, 232–244. 10.1002/evl3.28.

9 Narita, S., Kageyama, D., Hiroki, M., Sanpei, T., Hashimoto, S., Kamitoh, T., and Kato, Y. (2011). Wolbachia-induced feminisation newly found in Eurema hecabe, a sibling species of Eurema mandarina (Lepidoptera: Pieridae), Ecol. Entomol. 36, 309–317. 10.1111/j.1365-2311.2011.01274.x.

10 Hornett, E. A., Duplouy, A. M. R., Davies, N., Roderick, G. K., Wedell, N., Hurst, G. D. D., and Charlat, S. (2008). You can’t keep a good parasite down: evolution of a male-killer suppressor uncovers cytoplasmic incompatibility, Evolution 62, 1258–1263. 10.1111/j.1558-5646.2008.00353.x.

