## Supplementary material for "The rise and fall of feminizing *Wolbachia* in a butterfly": Document S1

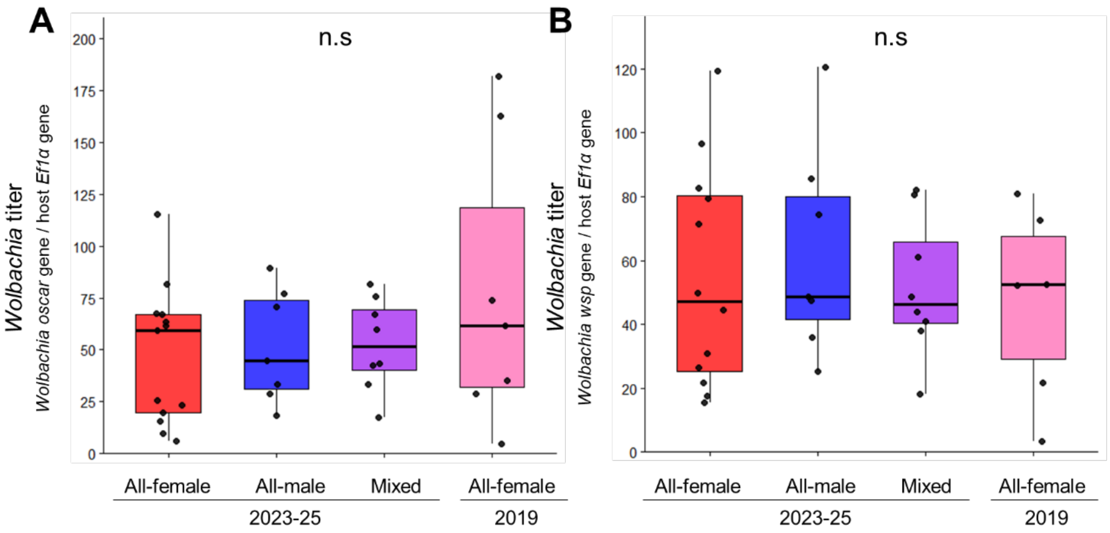


**Figure S1. *Wolbachia* titer in CF mothers producing offspring with different sex ratios.** Estimated *Wolbachia* densities in mothers that produced all-female broods, all-male broods, and mixed broods (i.e., broods with males and females) based on genomic qPCR. (A) Densities of *w*Fem estimated by *w*Fem-specific *oscar* gene copies per host nuclear *Ef1α* copy. (B) Densities of *w*CI estimated by *w*CI-specific *wsp* gene copies per host nuclear *Ef1α* copy.

**Table S1. Vertical transmission of *w*CI and *w*Fem from field-collected CF females to offspring.**

| ID of field-collected CF females | Offspring sex-ratio type | Offspring tested by PCR | Daughters | | Sons | |
|---|---|---|---|---|---|---|
| | | | CF | C | CF | C |
| 2023-IS222 | All-female | 4 daughters | 4 | 0 | – | – |
| 2023-IS226 | All-female | 10 daughters | 10 | 0 | – | – |
| 2023-IS258 | All-female | 13 daughters | 13 | 0 | – | – |
| 2023-IS227 | All-male | 7 sons | – | – | 0 | 7 |
| 2023-IS228 | All-male | 1 son | – | – | 0 | 1 |
| 2023-IS231 | All-male | 1 son | – | – | 0 | 1 |
| 2023-IS239 | Mixed | 1 daughter and 2 sons | 1 | 0 | 0 | 2 |
| 2023-IS251 | Mixed | 3 daughters and 4 sons | 3 | 0 | 0 | 4 |

CF, PCR-positive for *w*CI and *w*Fem. C, PCR-positive for *w*CI and PCR-negative for *w*Fem. –, not applicable.

### Supplemental experimental procedures

#### Field collection and laboratory rearing of *E. hecabe* butterflies

From 2023 to 2025, 640 *E. hecabe* adults were collected from 12 sites on Ishigaki Island, Japan. A total of 54 females collected between April 2023 and April 2025 were individually allowed to oviposit on the host plant *Ormocarpum cochinchinense* (Lour.) Merr. (Fabaceae). The collected eggs were separately reared in plastic containers until adult emergence under 16:8 h light-dark cycles at 25 °C. The hatched larvae were fed an artificial diet containing leaf powder of *Albizia julibrissin* Durazz. (Fabaceae), as described previously[S1]. The eggs and larvae were monitored daily. Eggs that did not hatch within 7 days were considered dead, regardless of any visible changes. The sex ratio of the emerged adults was recorded for each brood.

#### DNA extraction and diagnosis of *Wolbachia* infection in field-collected samples

Adult legs were homogenized in 100 µL of saline-Tris EDTA (STE) buffer (100 mM NaCl; 10 mM Tris-Cl [pH 8.0], and 1 mM EDTA) containing 2 µL of proteinase K (20 mg/mL). The homogenates were incubated at 37 °C for 30 min and then at 95 °C for 5 min. The supernatant obtained after centrifugation at 20,000 xg was used as the template for PCR.

The presence of *w*Fem was examined by PCR targeting the *oscar* gene using the forward primer Hm/Hb/Eh-Oscar.qF (5'–GGACCAGTAGAAGCACCTATATC–3')[S2] and reverse primer Hm/Hb/Eh-Oscar.qR (5'–AACACGACTAGCTTCAGGAC–3')[S2], which amplify a ca. 99 bp DNA fragment. The temperature profile adopted for PCR reaction was 94.0 °C for 1 min, 30 cycles of denaturation at 98 °C for 10 sec, annealing at 56 °C for 15 sec, and extension at 68 °C for 15 sec, followed by a final extension at 72 °C for 7 min. In addition to the feminizing *Wolbachia* strain *w*Fem, *E. hecabe* on Ishigaki Island is known to be fixed for *w*CI, which is not involved in sex ratio distortion. The presence of *w*CI was confirmed by PCR targeting the *wsp* gene using the *Wolbachia*-specific forward primer wsp81F (5'–TGGTCCAATAAGTGATGAAGAAAC–3')[S3] and the *w*CI-specific reverse primer WHecFem1 (5'–ACTAACGTTTTTGTTTAG–3')[S4], which amplify a 232-bp DNA fragment. PCR was performed for 35 cycles of denaturation at 95 °C for 1 min, annealing at 58 °C for 1.5 min, and extension at 72 °C for 1.5 min, followed by a final extension at 72 °C for 7 min.

To discriminate among the three *Eurema* species occurring on Ishigaki Island (*E. hecabe*, *E. mandarina*, and *E. blanda*), the extracted DNA was subjected to diagnostic PCR targeting the *Tpi* gene[S5]. *E. hecabe* was identified using the species-specific forward primer Eh6-F (5'–TGTGGCCTTCTGCCCTATTAAA–3') and reverse primer Eh6-R (5'–ACAGGCAATGACCTTGAGTC–3'), which amplify a 375-bp DNA fragment. The PCR conditions were 94.0 °C for 5 min, followed by 35 cycles of 94.0 °C for 30 s, 48 °C for 30 s, and 72.0 °C for

30 s, with a final extension at 72.0 °C for 7 min.

#### Inference of *Wolbachia* densities by quantitative real-time PCR

We estimated *Wolbachia* densities in mothers used in the 2023–2025 rearing experiments to determine whether the *Wolbachia* density differed among offspring sex-ratio types and whether it differed from that in CF mothers used in the 2019 rearing experiment[S6], which was conducted before the increase in the male frequency of the field population.

Thoracic tissues were collected from adult females sacrificed after oviposition and subjected to DNA extraction using the DNeasy Blood & Tissue Kit (Qiagen, Tokyo, Japan). The densities of *w*Fem and *w*CI were estimated by quantitative real-time PCR (qPCR), targeting the *Wolbachia oscar* gene for *w*Fem, the *wsp* gene for *w*CI, and the host nuclear *Ef1α* gene as an internal reference. *w*Fem and *w*CI were amplified using the same primer pairs as those used for the conventional PCR described above. The Ef1α gene was amplified using the forward primer EF-1F (5'–TCAGTTAAGGCTATTAACGCTCTG–3') and reverse primer EF-1R (5'–ATACTACCTTTTGCGGTTACTGTC–3')[S7]. The following cycling conditions were used for all primer pairs: 95 °C for 2 min, followed by 40 cycles of 95 °C for 5 s, 60 °C for 10 s, and 72 °C for 10 s.

*Wolbachia* titers were quantified using the ΔCt method, with the host *Ef1α* gene acting as the reference. Each sample was analyzed in technical duplicate. For statistical analysis, ΔCt values were analyzed using the Steel–Dwass test. For visualization, data were plotted as 2^−ΔCt values.

### Supplemental discussion

#### How can CF mothers produce male offspring? A possible underlying mechanism

Although C females have a ZW sex chromosome constitution, CF females are known to be Z0[S6,S8,S9]. For offspring produced by CF (Z0) mothers, four possible classes can be considered: Z0 individuals that inherit *w*Fem, Z0 individuals that fail to inherit *w*Fem, ZZ individuals that inherit *w*Fem, and ZZ individuals that fail to inherit *w*Fem.

The Z0 offspring that failed to inherit *w*Fem would be expected to develop as males because of the absence of the *w*Fem-derived protein Oscar[S10,S11]. However, such individuals are likely to be inviable because developing males carrying only one Z chromosome would suffer from inappropriate dosage compensation. Conversely, ZZ offspring that inherit *w*Fem would be feminized by *w*Fem-derived Oscar but are also expected to be inviable because the developing females carrying two Z chromosomes would suffer from inappropriate dosage compensation. Consequently, among the four possible classes, *w*Fem-lacking ZZ males and *w*Fem-carrying Z0 females are expected to survive. This is consistent with our observation that all sons produced by CF mothers lacked *w*Fem, whereas all daughters carried *w*Fem (Table S1).

We also examined the titers of *w*Fem and *w*CI in the CF mothers, but neither differed significantly among the mothers producing different offspring sex-ratio types (Steel–Dwass test, $P > 0.05$; Figure S1). This result suggests that the imperfect vertical transmission of *w*Fem may be caused by a failure of *w*Fem transmission within the ovaries, without a corresponding reduction in *w*Fem titer in somatic tissues, such as the thoracic tissue examined here. We hypothesize that a nuclear suppressor in CF mothers specifically targets *w*Fem in the germline, thereby inactivating the *w*Fem-induced prevention of Z chromosome inheritance to produce both Z0 and ZZ offspring[S9].

**Production of male offspring by CF mothers was also supported by the field data**

In our previous study, we analyzed mtDNA *COI* sequences of *E. hecabe* and found that all 20 CF females collected on Ishigaki Island carried haplotype 10, which was also shared by six C females[S12]. Thus, 76.9% (20 of 26) of females carrying haplotype 10 were infected with *w*Fem. In 2016, only one of 22 males (4.5%) carried haplotype 10. Because CF females were not known to produce male offspring at that time, this male was most plausibly derived from a C matriline carrying haplotype 10. In sharp contrast, 13 of 16 males (81.3%) collected in 2023 carried haplotype 10. This marked increase is consistent with the emergence of male production by CF females in 2023 and suggests that many of these males originated from CF matrilines. In 2025, the proportion of males carrying haplotype 10 declined to 58.3% (14 of 24), potentially reflecting the concurrent decline in *w*Fem prevalence in the population. Taken together, these findings provide field evidence consistent with our rearing experiments, suggesting that *w*Fem-infected mothers also produced sons under natural conditions, followed by a decline in *w*Fem prevalence in the population.

**Occurrence of *w*Fem-positive males among the field-collected males**

Among the field-collected males, 7 of 245 individuals were positive for *w*Fem (see Data availability), consistent with our previous detection of *w*Fem in 7 out of 88 individuals collected between 2019 and 2021[S6]. In contrast, no *w*Fem-positive males were obtained in our limited rearing experiments conducted in the present study: all offspring examined from the CF mothers were either CF females (n = 31) or C males (n = 15). The rarity of *w*Fem-positive males suggests that in most cases, maternal transmission of *w*Fem and *w*Fem-induced feminization are coupled. Nevertheless, the occurrence of occasional *w*Fem-positive males indicates that the threshold of *w*Fem density required to induce feminization would likely be higher than that required for maternal transmission, allowing *w*Fem to be transmitted to some offspring at a density insufficient to induce feminization.

**Acknowledgements**

We thank Masashi Nomura, Yuhei Hashimoto, and Rinka Kimoto for collecting the butterflies and Waka Minami for assistance with insect rearing. This work was supported by a JSPS KAKENHI grant (25K00108) and Sumitomo Foundation for basic Sciences to M.M. and a JSPS KAKENHI grant (23K26922 and 24H02293) to D.K.

**Author contributions**

K.I., D.K., and M.M. designed the study. K.I., A.M., and M.M. conducted field sampling, rearing experiments, and molecular experiments. K.I. and M.M. analyzed the data. K.I., D.K., and M.M. wrote the manuscript. All authors read and approved the final manuscript.

**Declaration of interests**

The authors declare no competing interests.

**Supplemental references**

S1. Kato, Y., and Sakakura, F. (1994). Artificial diet rearing of *Eurema blanda* and some notes on its host-plant, Trans. Lepid. Soc. Jpn. *45*, 21–26. 10.18984/lepid.45.1_21.

S2. Arai, H. Wijonarko, A., Katsuma, S., Naka, H., Kageyama, D., Hornett, E.A., Hurst, G.D.D. (2025) Evolution of *Wolbachia* male-killing mechanism within a host species, Curr. Biol. *35*, 2006–2018. 10.1016/j.cub.2025.03.027.

S3. Braig, H.R., Zhou, W.G., Dobson, S.L., and O’Neill, S.L. (1998). Cloning and characterization of a gene encoding the major surface protein of the bacterial endosymbiont *Wolbachia pipientis,* J. Bacteriol. *180*, 2373–2378. 10.1128/jb.180.9.2373-2378.1998.

S4. Hiroki, M., Tagami, Y., Miura, K., and Kato, Y. (2004). Multiple infection with *Wolbachia* inducing different reproductive manipulations in the butterfly *Eurema hecabe*, Proc. R. Soc. B *271*, 1751–1755. 10.1098/rspb.2004.2769.

S5. Miyata, M.N., Kageyama, D., and Nomura, M. (2020). Multiplex PCR for identification of two butterfly sister species: *Eurema mandarina* and *Eurema hecabe*, BMC Res. Notes *13*, 1–5. 10.1186/s13104-020-05093-3.

S6. Miyata, M., Nomura, M., and Kageyama, D. (2024). Rapid spread of a vertically transmitted symbiont induces drastic shifts in butterfly sex ratio, Curr. Biol. *34*, R490–R492. 10.1016/j.cub.2024.04.027.

S7. Kern, P., Cook, J.M., Kageyama, D., and Riegler, M. (2015). Double trouble: combined action of meiotic drive and *Wolbachia* feminization in *Eurema* butterflies, Biol. Lett. *11*, 20150095.

10.1098/rsbl.2015.0095.

S8. Narita, S., Kageyama, D., Hiroki, M., Sanpei, T., Hashimoto, S., Kamitoh, T., and Kato Y. (2011). *Wolbachia*-induced feminization newly found in *Eurema hecabe*, a sibling species of *Eurema mandarina* (Lepidoptera: Pieridae), Ecol. Entomol. *36*, 309–317. 10.1111/j.1365-2311.2011.01274.x.

S9. Kageyama, D., Ohno, M., Sasaki, T., Yoshido, A., Konagaya, T., Jouraku, A., Kuwazaki, S., Kanamori, H., Katayose, Y., Narita, S., et al. (2017). Feminizing *Wolbachia* endosymbiont disrupts maternal sex chromosome inheritance in a butterfly species, Evol. Lett. *1*, 232–244. 10.1002/evl3.28.

S10. Katsuma, S., Hirota, K., Matsuda-Imai, N., Fukui, T., Muro, T., Nishino, K., Kosako, H., Shoji, K., Takanashi, H., Fujii, T. et al. (2022). A *Wolbachia* factor for male killing in lepidopteran insects, Nat. Commun. *13*, 6764. 10.1038/s41467-022-34488-y.

S11. Arai, H., Nishikawa, Y., Konagaya, T., Kogawa, M., Kifushi, M., Takeyama, H., Anbutsu, H., Kageyama, D. (2025). Single-cell genome analysis of a feminizing *Wolbachia* in *Eurema* butterflies reveals a shared origin with male-killing *Wolbachia*, Microb. Genom. 11: 001578. 10.1099/mgen.0.001578.

S12. Miyata, M., Nomura, M., and Kageyama, D. (2025). Elusive origin of mitochondria shared by two sister species of *Eurema* butterflies, BMC Ecol. Evol. *25*, 71. 10.1186/s12862-025-02414-7.
